# IL-35 produced by dendritic cells via TIM-3-STAT3 signaling contributes to the development of visceral leishmaniasis

**DOI:** 10.64898/2026.02.23.707416

**Authors:** Sandeep Kumar, Shubham, Manish Mishra, Raj Kumar, Pradip Sen

**Author notes:** Department of Pathology, Yale School of Medicine, New Haven, CT, USA. These authors contributed equally to this work (Shubham, Manish Mishra).

## Abstract

Visceral leishmaniasis (VL), a life-threatening parasitic disease caused by *Leishmania donovani* (LD), progresses primarily due to profound immunosuppression. However, the molecular and cellular mechanisms underlying this immune dysfunction remain poorly defined. Here we show that early production of IL-35 by dendritic cells (DCs) is critical for immunosuppression and disease pathogenesis during LD infection. LD stimulated IL-35 expression in DCs through the TIM-3 receptor and the downstream transcription factor STAT3. IL-35 produced by DCs subsequently suppressed DC maturation and T cell proliferation, propagated immunosuppression by inducing IL-35 expression in T cells, and impaired type-1 anti-leishmanial immunity, thereby promoting disease progression. Genetic or pharmacologic inhibition of STAT3 markedly reduced IL-35 production by DCs, restored protective type-1 T cell responses, and promoted parasite clearance *in vivo*. Notably, treatment with WP1066, an FDA-designated orphan STAT3 inhibitor, significantly lowered parasite burden and disease severity in infected mice. Together, these findings uncover a previously unrecognized TIM-3-STAT3-IL-35 axis that drives immunosuppression and pathogenesis in VL and highlight STAT3 inhibition as a promising therapeutic strategy to restore host immunity and control infection.

## INTRODUCTION

Visceral leishmaniasis (VL), a potentially fatal disease caused by *Leishmania donovani* (LD), has remained severely neglected, even though the WHO recognizes it as the world’s second most important parasitic infection (Rostan *et al*, 2013). Alarmingly, VL is now spreading to non-endemic regions, including developed countries such as the United States, the United Kingdom, and parts of Europe (Curtin & Aronson, 2021; Donnelly *et al*, 2025; Mody *et al*, 2019; Pasquier *et al*, 2022; Shrestha *et al*, 2019; Stahlman *et al*, 2017), raising concerns about its expanding global footprint and the emergence of a new public health threat in these areas. The pathogenesis of VL is primarily driven by profound immunosuppression (Flora *et al*, 2014). However, the cellular and molecular mechanisms underlying this immunosuppression remain largely undefined. To date, only interleukin (IL)-10 and transforming growth factor-beta (TGF-β) are known to suppress host immunity during LD infection (Akhtar *et al*, 2022; Bunn *et al*, 2018; Mishra *et al*, 2023; Nylen & Sacks, 2007; Rodrigues *et al*, 1998; Yadav *et al*, 2023), leaving open the possibility that other cytokines may also play immunosuppressive roles in promoting VL pathogenesis.

In recent years, IL-35 has emerged as an important immunosuppressive cytokine (Sawant *et al*, 2015). IL-35 belongs to the IL-12 cytokine family, which includes three other cytokines: IL-12, IL-23 and IL-27 (Chen *et al*, 2018). It is a heterodimeric cytokine composed of the p35 subunit of IL-12 (encoded by *IL12A*) and the Epstein-Barr virus-induced gene 3 subunit (EBI3, encoded by *EBI3*) of IL-27 (Turnis *et al*, 2016; Vignali & Kuchroo, 2012). IL-35 was first discovered to be constitutively produced by regulatory T (T_reg_) cells and to inhibit the proliferation of effector T (T_eff_) cells (Vignali & Kuchroo, 2012). Subsequent studies have shown that IL-35 can also be produced by other cells, such as dendritic cells (DCs). However, evidences for IL-35 production by DCs are conflicting. While one study demonstrated IL-35 expression in dexamethasone-treated human DCs (Dixon *et al*, 2015), another failed to detect IL-35 expression in human DCs (Chen *et al*., 2018). Given this contradiction, it is still unclear whether DCs express IL-35 at all.

The role of IL-35 in VL is similarly obscure. Although a recent study reported that elevated expression of the IL-35 subunit EBI3 in CD4^+^ T cells contributes to disease progression (Asad *et al*, 2019), that study focused solely on EBI3 and did not examine the role of the complete IL-35 heterodimer. Because EBI3 itself can exert immunosuppressive effects (Hildenbrand *et al*, 2023), the precise contribution of IL-35 to VL pathogenesis remains unclear. In fact, it has not been established whether the IL-35 heterodimer is produced during LD infection. Moreover, if IL-35 is expressed during LD infection, its initial cellular source must be defined to understand how early IL-35 production shapes T cell responses, anti-leishmanial immunity, and disease pathogenesis. Considering that DCs become infected by *Leishmania* parasites and play a central role in initiating anti-leishmanial T cell responses (Bennett *et al*, 2001; Martin *et al*, 2010), we examined whether they serve as early producers of IL-35. We also sought to delineate the signaling pathway responsible for IL-35 induction in DCs, as its regulation remains poorly defined. Our results show that DCs produce IL-35 upon LD infection and act as the earliest cellular source of this cytokine. Mechanistically, LD induces IL-35 expression in DCs through a pathway involving the inhibitory receptor TIM-3 (T cell immunoglobulin and mucin domain-containing protein-3) and its downstream effector, the STAT3 transcription factor, neither of which had previously been shown to regulate IL-35 expression. DC-derived IL-35 subsequently promotes IL-35 expression in T cells, suppresses DC maturation and T cell proliferation, and attenuates anti-leishmanial immunity, thereby facilitating disease progression. Notably, pharmacological inhibition of the TIM-3-STAT3 pathway using WP1066, an FDA-approved STAT3 inhibitor for the treatment of pediatric brain tumors (Wang *et al*, 2022) (https://www.moleculin.com/technology/wp1066/), significantly reduces IL-35 expression in DCs and lowers parasite burden *in vivo*. Together, these findings establish DC-derived IL-35 as a key immunosuppressive mediator of VL pathogenesis, identify the TIM-3-STAT3 axis as a novel upstream regulator of IL-35 expression, and suggest WP1066 as a potential therapeutic for VL.

## RESULTS

### DCs produce IL-35 during LD infection

Because it was previously unknown whether DCs produce IL-35 in response to LD infection, we first investigated this aspect. We infected bone marrow-derived DCs (BMDCs) with LD promastigotes (LDPm; extracellular form) for varying durations and assessed IL-35 expression by flow cytometric detection of cells coexpressing EBI3 and IL-12p35 (EBI3^+^IL-12p35^+^). The frequency of EBI3^+^IL-12p35^+^ BMDCs increased as early as 6 h post-LDPm infection, peaked at 48 h, and remained elevated for at least 60 h (Fig. 1A). Similar to LDPm, infection with LD amastigotes (LDAm; intracellular form) also increased the frequency of EBI3^+^IL-12p35^+^ BMDCs (Fig. 1B). Consistent with these protein-level changes, reverse transcription quantitative PCR (RT-qPCR) analysis revealed significantly increased expression of *EBI3* and *IL12A* mRNAs in BMDCs infected with LDPm for 12 or 24 h compared with uninfected controls (Fig. 1C). Because EBI3 and IL-12p35 can also pair with alternative subunits to form IL-27 and IL-12, respectively (Sawant *et al*., 2015), coexpression alone does not establish formation of the IL-35 heterodimer. We therefore directly examined whether these subunits associate to form the IL-35 complex in DCs following LD infection. Confocal microscopy revealed enhanced intracellular colocalization of EBI3 and IL-12p35 in LDPm-infected BMDCs compared with uninfected cells (Fig. 1D). This interaction was further confirmed biochemically by co-immunoprecipitation and spatially by confocal fluorescence resonance energy transfer (c-FRET), demonstrating that EBI3 and IL-12p35 indeed form the IL-35 heterodimer in LD-infected BMDCs (Fig. 1, E and F). The increased IL-35 complex formation in LD-infected BMDCs likely results from the elevated expression of EBI3 and IL-12p35 in these cells compared to uninfected BMDCs (Fig. 1E). To determine whether DCs secrete IL-35 upon LD infection, we measured IL-35 in culture supernatants using IL-35 heterodimer-specific ELISA. LDPm-infected BMDCs secreted significantly higher amounts of IL-35 than uninfected controls (Fig. 1G), establishing that LD infection induces IL-35 production by DCs *in vitro*. Notably, LD infection similarly increased IL-35 expression in human monocyte-derived DCs (HuMoDCs), indicating that this response is conserved across species (Fig. 1H). Next, we examined whether DCs produce IL-35 during LD infection *in vivo*. Accordingly, we isolated splenocytes from LD-infected mice at various time points postinfection and analyzed IL-35 expression in splenic DCs (sDCs), identified by CD11c^+^ cells (Chiba *et al*, 2012; Ogawa *et al*, 2014; Stager *et al*, 2006; Wang *et al*, 2021; Wu *et al*, 2018), by flow cytometry. The frequency of IL-35-expressing sDCs increased by day 22 postinfection and continued to rise through days 30 and 60 (Fig. 1I, red dots in the right panel; and Fig. EV1A), demonstrating sustained IL-35 production by DCs during LD infection. In contrast, IL-35 expression in splenic T cells and other non-DC, non-T cell populations (CD11c^-^CD3^-^ cells, indicated as “other cells”) was minimal at early time points and increased only modestly at later stages of infection (Fig. 1I, green and blue dots in the right panel; and Fig. EV1A). Moreover, DCs consistently expressed higher levels of the IL-35 subunits EBI3 and IL-12p35 than other splenocyte populations throughout the course of infection (Fig. EV1B). Together, these findings establish that LD infection induces DCs to produce the IL-35 heterodimer both *in vitro* and *in vivo*, with DCs serving as the earliest and predominant cellular source of IL-35 during infection.

**Figure 1.**
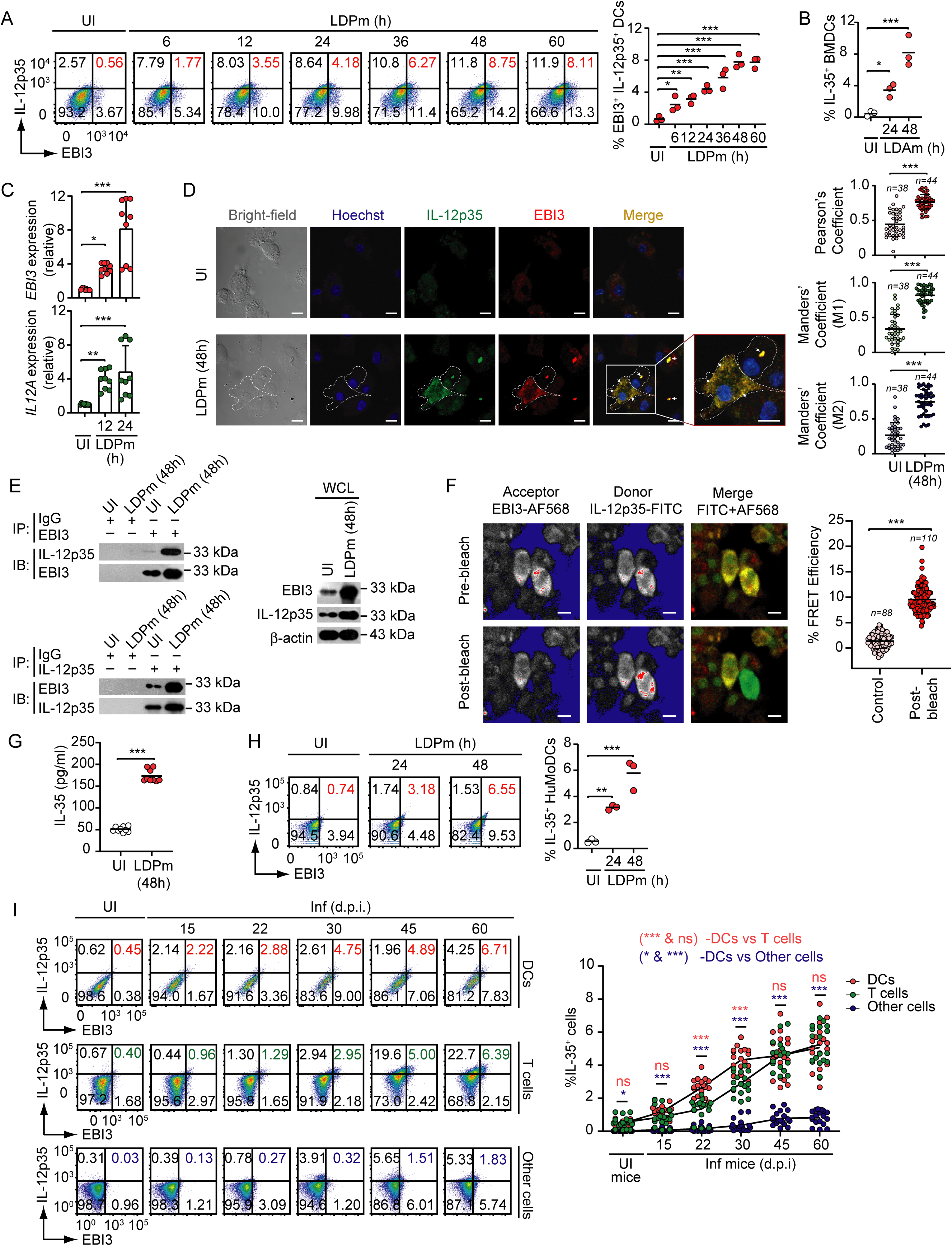
LD induces IL-35 production in DCs. (**A**) The frequency of IL-35-expressing (i.e. IL-12p35^+^EBI3^+^) BMDCs, either uninfected (UI) or infected with LDPm for indicated time points, was determined by flow cytometry. In this and other flow cytometry figures, numbers in each quadrant indicate the percentage of cells in the respective quadrant (representative of *n* = 3 experiments; left). Right: summary of three experiments. (**B**) The frequency of IL-35 expressing BMDCs infected with LDAm for indicated time points was analyzed by flow cytometry as described in (A) and is presented graphically (data pooled from three experiments). (**C**) *EBI3* and *IL12A* mRNA expression in uninfected BMDCs and BMDCs infected with LDPm for 12 or 24 h was assessed by RT-qPCR. Results were normalized to *ACTB* mRNA (encoding β-actin) expression and are presented as fold change relative to uninfected BMDCs (*n* = 9 replicates per group). (**D**) Confocal microscopic analysis of the colocalization (merge; yellow) of IL-12p35 (green) and EBI3 (red) in uninfected and LDPm-infected (48 h) BMDCs; nuclei were stained with Hoechst (blue) (representative of *n* = 3 experiments; left). Scale bar, 10 μm. Right: IL-12p35/EBI3 colocalization quantified by Pearson’s and Manders’ Coefficients. (**E**) The association between IL-12p35 and EBI3 in uninfected BMDCs or BMDCs infected with LDPm for 48 h was assessed by immunoprecipitation (IP) followed by immunoblotting (IB); β-actin serves as a loading control (representative of *n* = 3 experiments). WCL, whole-cell lysate (no IP); IgG, immunoglobulin G (IP control). (**F**) Interaction between EBI3 and IL-12p35 in BMDCs infected with LDPm for 48 h, assessed by FRET (representative of *n* = 3 experiments; left). Scale bar, 10 μm. Right: FRET efficiency. (**G**) IL-35 production by uninfected and LDPm-infected (48 h) BMDCs measured by ELISA (combined data from three experiments, each with *n* = 3 replicates). (**H**) HuMoDCs were infected with LDPm for indicated times, and the frequency of IL-35-expressing DCs was analyzed by flow cytometry as in (A) (representative plots from *n* = 3 experiments; left). Right: pooled data from three independent experiments. (**I**) Frequency of IL-35-expressing DCs, T cells, and other cells (i.e., non-DC, non-T cells; CD11c^-^CD3^-^ cells) in the spleen of LD-infected mice at indicated days postinfection, analyzed by flow cytometry [representative plots (left) and pooled data (right); *n* = 18 mice per time point]. The gating strategy is shown in Fig. EV1A. The levels of the IL-35 subunits EBI3 and IL-12p35 in these cell populations is shown in Fig. EV1B. Each symbol represents data from one experiment [A (right panel), B and H (right panel)], replicate (C and G), field [D (right panel)], cell [F (right panel)], or mouse [I (right panel)]. Horizontal bars (B, G, and right panels of A, D, F and H) indicate means and error bars (C, D and F) represent SD. \**P* < 0.05, \*\**P* < 0.01, \*\*\**P* < 0.001; ns, not significant.

### Regulation of LD-induced IL-35 expression in DCs

We then investigated how LD enhances IL-35 production in DCs. In this context, we first explored the role of the transcription factor STAT3 in this process. This is because our recent findings (Mishra *et al*., 2023) showed that the transcription factor STAT3 becomes activated in DCs during LD infection, with activation peaking at 0.5 h and persisting up to 6 h postinfection. Additionally, a previous report (Collison *et al*, 2012) indicated the presence of potential STAT-binding sites in the promoters of IL-35 subunit: ^-210^TTCTAGAA^-203^ and ^-386^TTCCCAGAGA^-377^ in the *EBI3* promoter, and ^-113^TCCTGGGAA^-105^, ^-365^TTCTTTAGAA^-356^, ^-1567^CTCTAGGAAA^-1558^ and ^-1702^TTCTACAAGAA^-1692^ in the *IL12A* promoter [the base positions are relative to the transcription start sites (Liu *et al*, 2003; Wirtz *et al*, 2005)] (Fig. 2A, top). However, it remains to be confirmed whether the STAT transcription factors actually bind to these sites. Moreover, it is not yet known whether STAT3 can bind to the *EBI3* and *IL12A* promoters, and if so, whether LD infection can trigger such binding. To address these questions, we performed chromatin immunoprecipitation (ChIP) analysis using BMDCs that were either uninfected or infected with LDPm for 0.5 h, a time point corresponding to maximal STAT3 activation (Mishra *et al*., 2023). Our results showed that LDPm induced STAT3 recruitment to the *EBI3* promoter regions -249/-120 and -436/-328 and to the *IL12A* promoter region -415/-306 in DCs, which contain the putative STAT-binding sites ^-210^TTCTAGAA^-203^, ^-386^TTCCCAGAGA^-377^ and ^-365^TTCTTTAGAA^-356^, respectively (Fig. 2A, bottom). To directly test whether these sites bind STAT3 upon LD infection, we performed DNA pull-down assay. We found that the *EBI3* and *IL12A* promoter-specific oligonucleotides EBI3-Pr1, EBI3-Pr2 and IL12A-Pr containing the wild-type STAT-binding sites (^-210^TTCTAGAA^-203^,^-386^TTCCCAGAGA^-377^ and ^-365^TTCTTTAGAA^-356^, respectively) precipitated STAT3 from nuclear lysates of LD-infected DCs (Fig. 2B). In contrast, the same oligonucleotides carrying mutations at these STAT-binding sites (MutEBI3-Pr1, MutEBI3-Pr2 and MutIL12A-Pr) failed to precipitate STAT3 (Fig. 2B). These results indicate that ^-210^TTCTAGAA^-203^ and ^-386^TTCCCAGAGA^-377^ in the *EBI3* promoter, and ^-365^TTCTTTAGAA^-356^ in the *IL12A* promoter, function as STAT3-binding sites.

**Figure 2.**
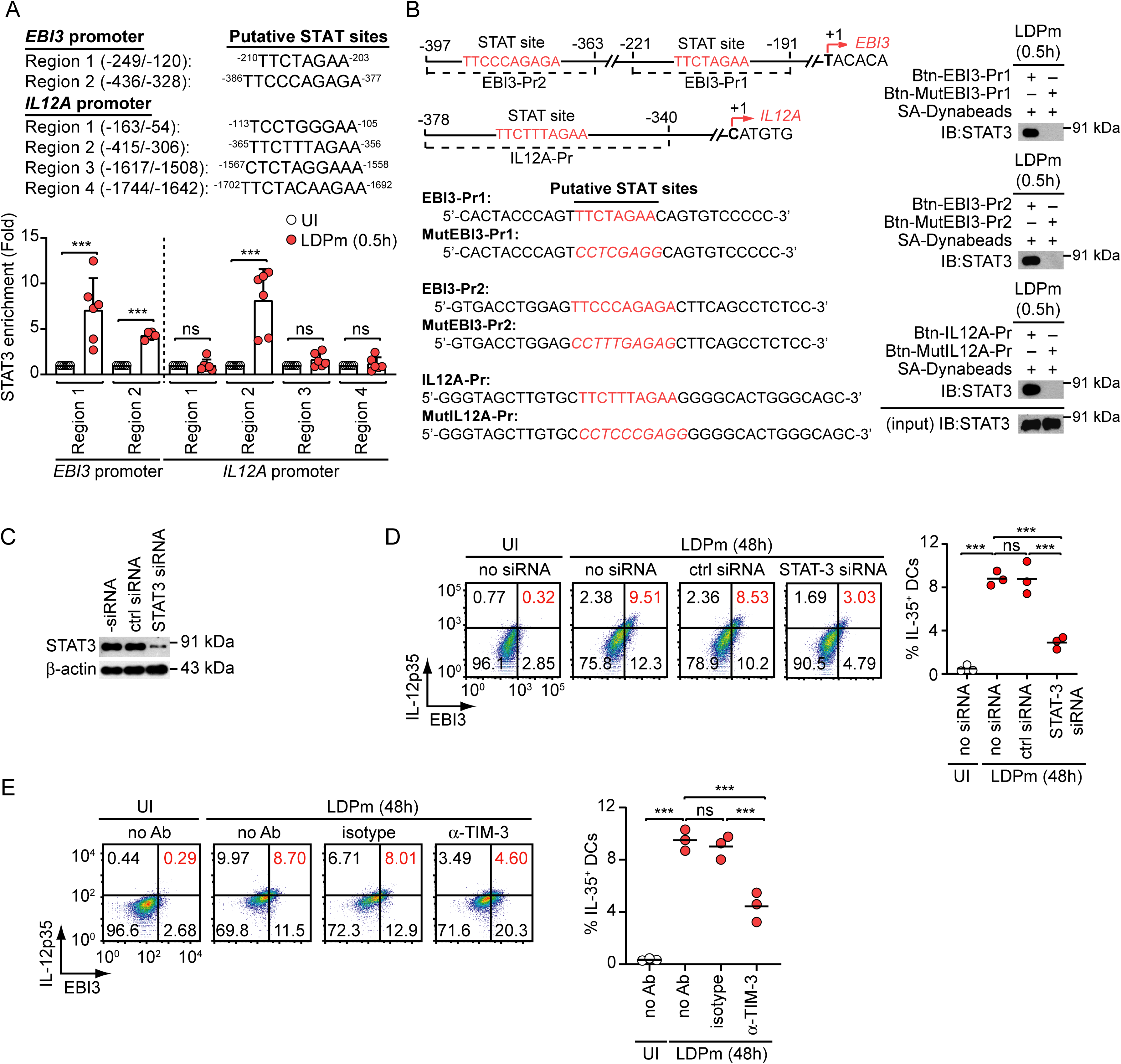
Role of TIM-3 and STAT3 in LD-induced IL-35 production in DCs. (**A**) Top: putative STAT sites in the *EBI3* and *IL12A* promoters. Bottom: ChIP-qPCR analysis of STAT3 recruitment to the indicated regions of *EBI3* and *IL12A* promoters in BMDCs at 0.5 h after LDPm infection (*n* = 6 replicates). Results are presented as fold enrichment relative to uninfected BMDCs. (**B**) Left: Details of *EBI3* and *IL12A* promoter-specific oligonucleotides containing wild-type or mutated STAT sites (mutated bases in italics) used for the DNA pull-down assay. Right: DNA pull-down analysis using streptavidin (SA)-conjugated Dynabeads, followed by immunoblotting to assess the binding of STAT3 [present in the nuclear lysates of LDPm-infected (0.5 h) BMDCs] to the biotin (Btn)-labeled oligonucleotides shown in the left panel (representative of *n* = 3 experiments). (**C**) Immunoblot analysis confirming STAT3 silencing by siRNA; β-actin serves as a loading control (representative of *n* = 3 experiments). Ctrl siRNA, control siRNA. (**D**) IL-35 expression in uninfected and LDPm-infected BMDCs (48 h infection) transfected with the indicated siRNAs, analyzed by flow cytometry [representative data (left) and compiled data (right) from *n* = 3 experiments]. (**E**) Effect of TIM-3 blockade using an anti-TIM-3 antibody on IL-35 production by BMDCs infected with LDPm for 48 h, analyzed by flow cytometry [representative (left) and compiled (right) data from *n* = 3 experiments]. Uninfected BMDCs without any antibody treatment (no Ab) serve as controls. Each symbol represents data from one replicate (A) or one experiment (right panels of D and E). Horizontal bars (right panels of D and E) denote means; error bars (A) indicate SD. \*\*\**P* < 0.001; ns, not significant.

We next performed reporter assays to determine whether STAT3 binding to these sites activates the *EBI3* and *IL12A* promoters. Overexpression of STAT3 in HEK293T cells increased wild-type *EBI3* and *IL12A* promoter activity, whereas mutation of the STAT3-binding sites significantly attenuated this effect (Fig. EV2, A and B). Consistent with these findings, silencing of STAT3 expression by small interfering RNA (siRNA) inhibited LDPm-induced IL-35 expression in BMDCs (Fig. 2, C and D). Thus, STAT3 enhances IL-35 expression by activating the *EBI3* and *IL12A* promoters.

Our recent study demonstrated that LD induces STAT3 activation through the TIM-3 receptor (Mishra *et al*., 2023). Accordingly, we examined the role of TIM-3 in LD-induced IL-35 production in DCs by blocking TIM-3 using an anti-TIM-3 antibody (Akhtar *et al*., 2022). Whereas LDPm readily induced IL-35 production in BMDCs treated with isotype control antibody, blockade of TIM-3 with anti-TIM-3 antibody reduced this effect (Fig. 2E). Collectively, these results demonstrate that LD induces IL-35 production in DCs via the TIM-3 receptor and its downstream effector STAT3. Furthermore, STAT3 mediates this effect by activating the *IL12A* and *EBI3* promoters.

### DC-derived IL-35 suppresses DC and T cell responses and impairs anti-leishmanial immunity

Since our findings indicated that DCs are early producers of IL-35 during LD infection (Fig. 1I), we next examined the potential effects of DC-secreted IL-35 on bystander DCs and T cells. For this, we incubated BMDCs (from uninfected mice) for 24 h with culture supernatants from uninfected or LD-infected DCs in the presence or absence of isotype control antibody or neutralizing anti-IL-35 antibody (clone V1.4C4.22) (Collison *et al*, 2010). We then stimulated BMDCs with lipopolysaccharide (LPS) for 24 h and assessed DC activation and maturation by analyzing the expression of co-stimulatory molecules (CD40, CD80 and CD86) and the secretion of pro-inflammatory cytokines [IL-12p70 and tumor necrosis factor alpha (TNFα)] via flow cytometry and ELISA, respectively. Preincubation with LD-infected DC supernatants, but not with uninfected DC supernatants, significantly suppressed LPS-induced upregulation of co-stimulatory molecule expression and pro-inflammatory cytokine secretion by BMDCs (Fig. 3, A and B). Treatment with anti-IL-35 antibody, however, reversed these inhibitory effects (Fig. 3, A and B), demonstrating that LD-infected DC supernatants impair DC activation and maturation in an IL-35-dependent manner.

**Figure 3.**
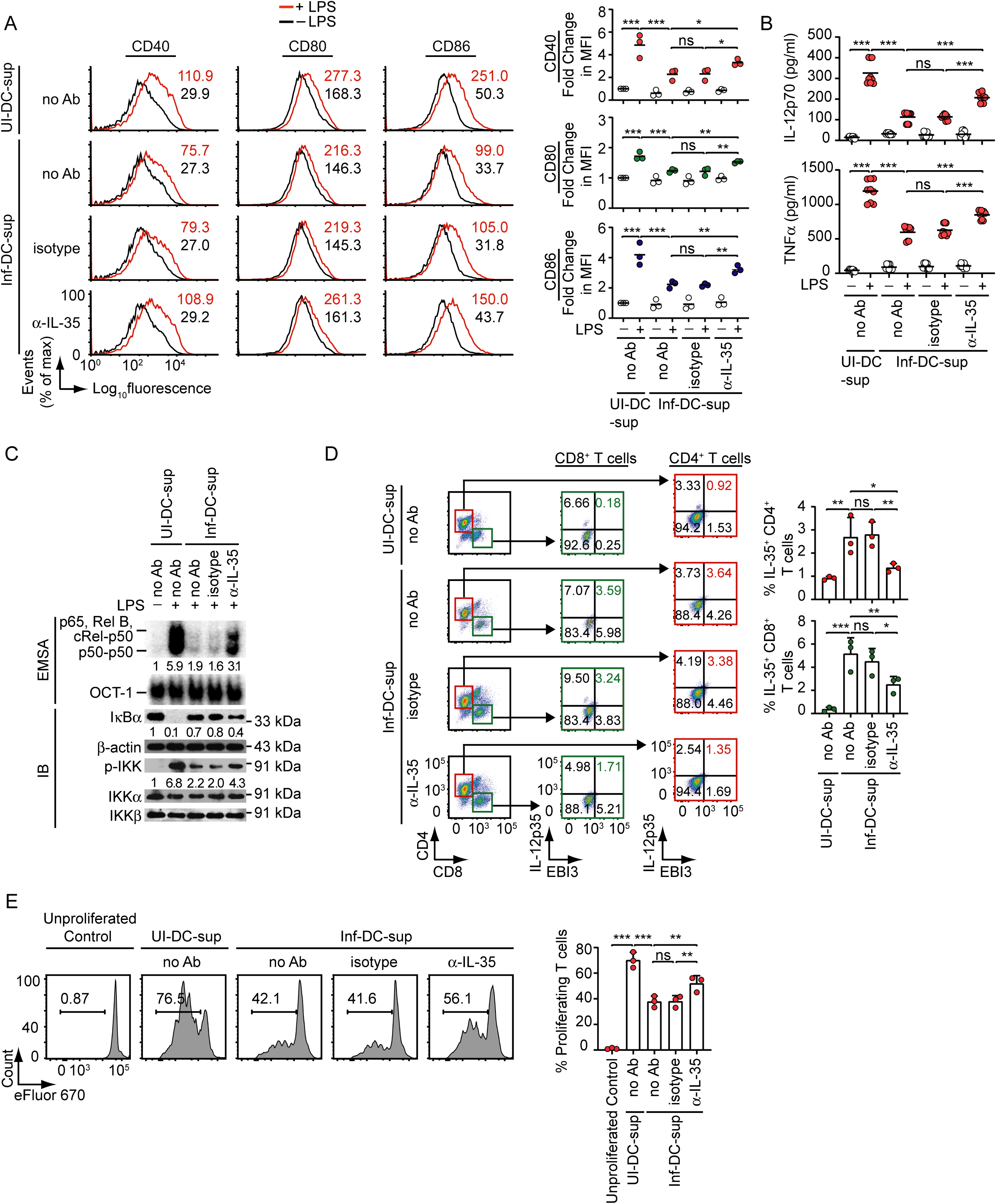
IL-35 from LD-infected DCs inhibits DCs and T cell activation. (**A** to **C**) BMDCs from uninfected mice were treated for 24 h with supernatants from uninfected DCs (UI-DC-sup) or LDPm-infected DCs (Inf-DC-sup), together with the indicated antibodies. Cells were then stimulated with LPS for an additional 24 h (A and B) or 0.5 h (C). The expression of costimulatory molecules on BMDCs was analyzed by flow cytometry [A, left: representative histograms with numbers indicating MFIs (isotype control-subtracted) and right: combined relative MFI data (presented as fold change over BMDCs treated with UI-DC-sup without antibody and LPS) from three experiments]. Cytokine secretion (IL-12 and TNFα) by BMDCs was measured by ELISA (B, *n* = 9 replicates). Nuclear NF-κB and OCT-1 (internal control) DNA-binding activities were examined by EMSA, while IκBα and phosphorylated (p-) IKK levels were assessed by immunoblotting using β-actin, IKKα and IKKβ as loading controls (**C**, representative of *n* = 3 experiments). Densitometry data (C, below lanes) were normalized to OCT-1 binding for EMSA, to β-actin expression for IκBα, and to the mean of IKKα and IKKβ expression for p-IKK in immunoblot analyses (IB). See Fig. EV3 for densitometry data of *n* = 3 experiments. (**D** and **E**) T cells from uninfected mice were treated with indicated supernatants and antibodies for 72 h in anti-CD3/anti-CD28-coated plates. In (E) T cells were labeled with eFluor 670 dye prior to these treatments. IL-35 expression in CD8^+^ and CD4^+^ T cells (D) and T cell proliferation (E) were analyzed by flow cytometry [representative data (left) and combined data (right) from *n* = 3 experiments]. In [E (left panel)], numbers above bracketed lines indicate the percentage of proliferating T cells. Each symbol represents data from a single experiment (the right panels of A, D and E), or a replicate (B). Horizontal bars in [A (right panel)] and (B) indicate mean values, and error bars in (D) and (E) represent SD. \**P* < 0.05, \*\**P* < 0.01, \*\*\**P* < 0.001; ns, not significant.

Because DC activation and maturation are primarily driven by nuclear factor (NF)-κB signaling (Sen *et al*, 2007), we subsequently investigated whether IL-35 in LD-infected DC supernatants inhibits DCs by suppressing the NF-κB pathway. While LD-infected DC supernatants inhibited LPS-stimulated NF-κB DNA-binding activity, IκBα degradation and IKK phosphorylation in BMDCs, the addition of a neutralizing anti-IL-35 antibody negated these effects (Fig. 3C and Fig. EV3). These findings indicate that IL-35 produced by DCs during LD infection suppresses the activation and maturation of bystander DCs by inhibiting NF-κB signaling. We then assessed whether IL-35-producing LD-infected DCs develop a tolerogenic phenotype. To address this, we infected DCs with LDPm and analyzed the expression of the tolerogenic marker indoleamine 2,3-dioxygenase (IDO) (Jimenez-Leon *et al*, 2023) in IL-35-producing DCs by flow cytometry. We found that LDPm infection considerably increased IDO expression in IL-35-producing BMDCs (Fig. EV4A). Consistent with these findings, LD-infected mice showed a higher percentage of IL-35^+^ sDCs expressing IDO compared with uninfected mice at all postinfection time points (Fig. EV4B). Therefore, DCs that produce IL-35 during LD infection not only inhibit the activation and maturation of bystander DCs but also exhibit a tolerogenic phenotype.

Next, we investigated whether DC-derived IL-35 exerts any effect on T cells. Accordingly, we examined whether IL-35 produced by DCs influences IL-35 expression in T cells. To test this, we cultured splenic T cells from uninfected mice with uninfected or LD-infected DC supernatants in the presence of an isotype control antibody or a neutralizing anti-IL-35 antibody and assessed IL-35 expression in CD4^+^ and CD8^+^ T cells via flow cytometry. LD-infected DC supernatants, but not uninfected DC supernatants, increased IL-35 expression in both CD4^+^ and CD8^+^ T cells; however, the addition of anti-IL-35 antibody blocked this effect (Fig. 3D). Thus, IL-35 secreted by LD-infected DCs promotes IL-35 expression in T cells.

We then determined whether IL-35 from LD-infected DCs influences T cell activation. For this, we isolated splenic T cells from uninfected mice, labeled them with eFluor 670 dye, and activated them with anti-CD3 and anti-CD28 antibodies in the presence or absence of uninfected or LD-infected DC culture supernatants together with anti-IL-35 or isotype control antibody. We measured T cell proliferation by flow cytometry. Compared with uninfected DC supernatants, LD-infected DC supernatants inhibited T cell proliferation despite anti-CD3 and anti-CD28 stimulation (Fig. 3E). Notably, addition of the neutralizing antibody substantially reduced this inhibitory effect (Fig. 3E). These findings indicate that IL-35 produced by LD-infected DCs inhibits T cell proliferation.

Having established that DC-derived IL-35 suppresses DC and T cell responses *in vitro*, we next examined whether IL-35 produced by DCs impairs anti-leishmanial immunity *in vivo*. Because DC-restricted genetic targeting of IL-35 is not currently available, we adopted an alternative strategy to selectively neutralize IL-35 produced by DCs during infection. Specifically, we transfected BMDCs with a neutralizing anti-IL-35 antibody and adoptively transferred these cells into syngeneic LD-infected mice at defined time points postinfection (Fig. 4A), with the aim of neutralizing IL-35 produced within the transferred DCs during their interaction with the parasite *in vivo*. We confirmed the effectiveness of this approach by demonstrating that the transfected anti-IL-35 antibody efficiently bound intracellular IL-35 in IL-35-overexpressing BMDCs (Fig. 5, A to C). In addition, previous reports demonstrating that adoptively transferred BMDCs persist only transiently *in vivo*, yet effectively modulate T cell priming and differentiation during early immune interactions, further support the suitability of this DC adoptive transfer strategy (Huang *et al*, 2013; Mishra *et al*., 2023; Schimmelpfennig *et al*, 2005; Xu *et al*, 2020). On day 60 postinfection, we assessed disease progression by measuring spleen and liver weights, parasite burden, and the frequencies of splenic CD4⁺ and CD8⁺ T cells producing the host-protective type-1 cytokine IFN-γ (Akhtar *et al*, 2020) or the disease-promoting type-2 cytokine IL-10 (Mishra *et al*., 2023). Additionally, we measured the frequency of splenic T cells expressing IL-35. Unlike control antibody-transfected BMDCs, adoptive transfer of anti-IL-35 antibody-transfected BMDCs reduced hepatosplenomegaly and parasite burden in the liver and spleen of LD-infected mice (Fig. 4, B and C). Furthermore, adoptive transfer of BMDCs transfected with anti-IL-35 antibody increased the frequency of IFNγ-expressing CD4^+^ and CD8^+^ T cells and decreased the frequency of IL-10-expressing CD4^+^ and CD8^+^ T cells in the spleens of LD-infected mice (Fig. 4D and Fig. 6A). These findings mirror the effects of *in vivo* IL-35 neutralization, which similarly promoted a protective T cell response and reduced parasite burden in LD-infected mice (Fig. EV7, A to D). Importantly, adoptive transfer of anti-IL-35 antibody-transfected BMDCs also decreased the frequency of IL-35-expressing T cells in the spleen of LD-infected mice (Fig. 4E and Fig. EV6B), providing *in vivo* evidence that DC-derived IL-35 drives IL-35 expression in T cells during LD infection. Collectively, these findings suggest that IL-35 produced by DCs during LD infection impedes DC and T cell activation, drives IL-35 expression in T cells, and suppresses anti-leishmanial immune responses, thereby facilitating disease progression.

**Figure 4.**
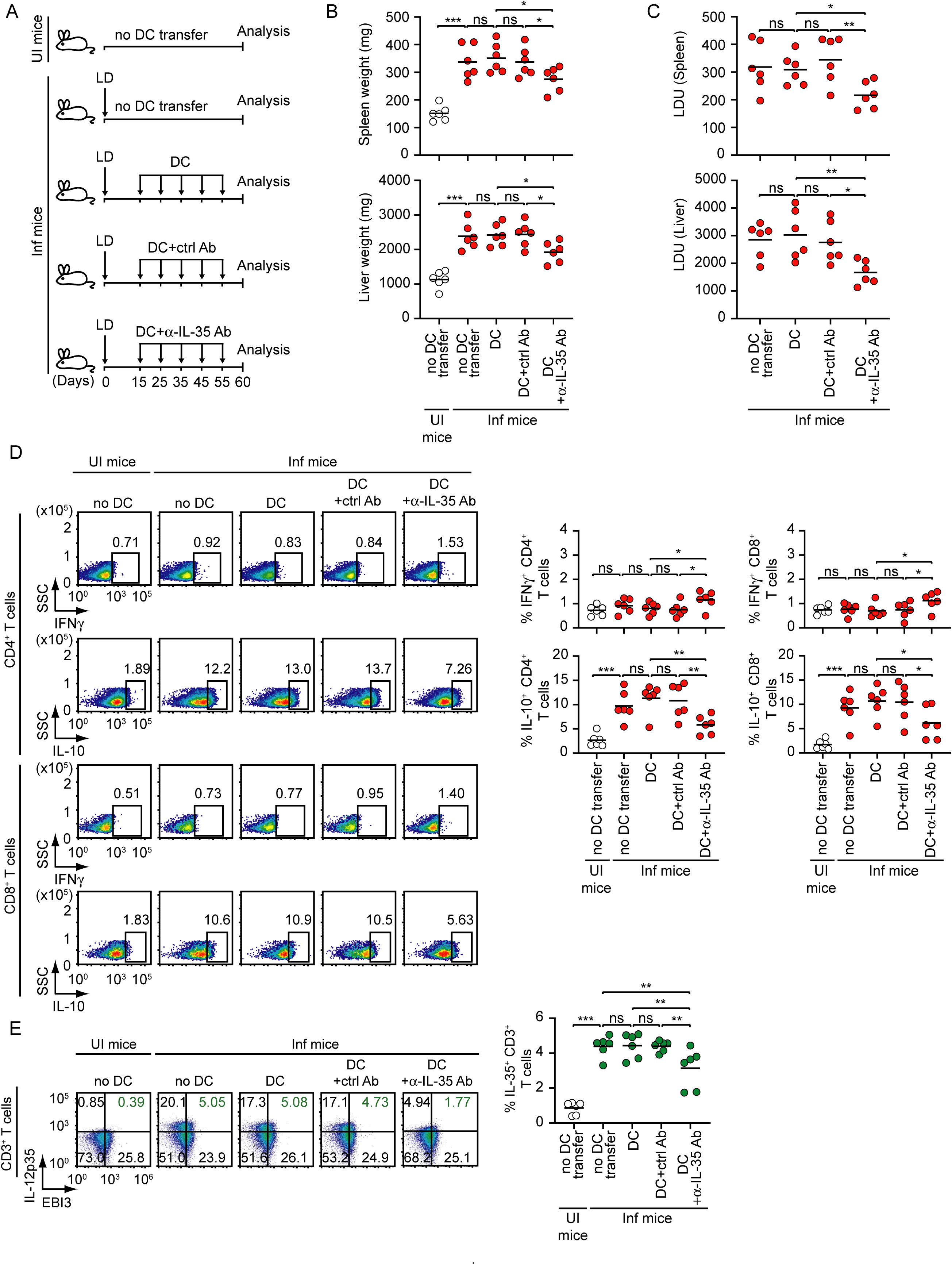
DC-derived IL-35 inhibits anti-leishmanial T cell response and promotes disease pathogenesis. (**A**) Schematic of the adoptive transfer protocol for anti-IL-35 antibody-transfected DCs: BALB/c BMDCs (1 x 10^6^), either untransfected or transfected with an isotype control (Ctrl) or a neutralizing anti-IL-35 antibody (transfection efficiency shown in Fig. EV5B), were adoptively transferred intravenously into LD-infected BALB/c mice on the indicated days postinfection (shown by arrows). In some experiments, no DC was transferred into LD-infected or uninfected mice. At day 60 postinfection, spleen and liver of these mice were collected for subsequent analyses (see panels B to E). (**B** and **C**) Spleen and liver weights (B) and parasite burdens (C; expressed as LDU) are shown (combined data from two experiments; *n* = 3 mice per group in each experiment). (**D** and **E)** Frequencies of IFNγ- or IL-10-producing CD4^+^ and CD8^+^ T cells (D) and IL-35-expressing total T cells (IL-12p35^+^EBI3^+^ CD3^+^ cells; E) in the spleen were analyzed by flow cytometry. Numbers above the outlined regions (D) or within quadrants (E) indicate the percentage of cells in the respective region or quadrant (representative of *n* = 6; left). Right: combined data from two separate experiments (*n* = 3 mice per group in each experiment). Gating strategies are shown in Fig. EV6, A and B. In (B), (C), and the right panels of (D) and (E), each symbol represents the data from one mouse, and horizontal bars indicate mean values. \**P* < 0.05, \*\**P* < 0.01, \*\*\**P* < 0.001; ns, not significant.

**Figure 5.**
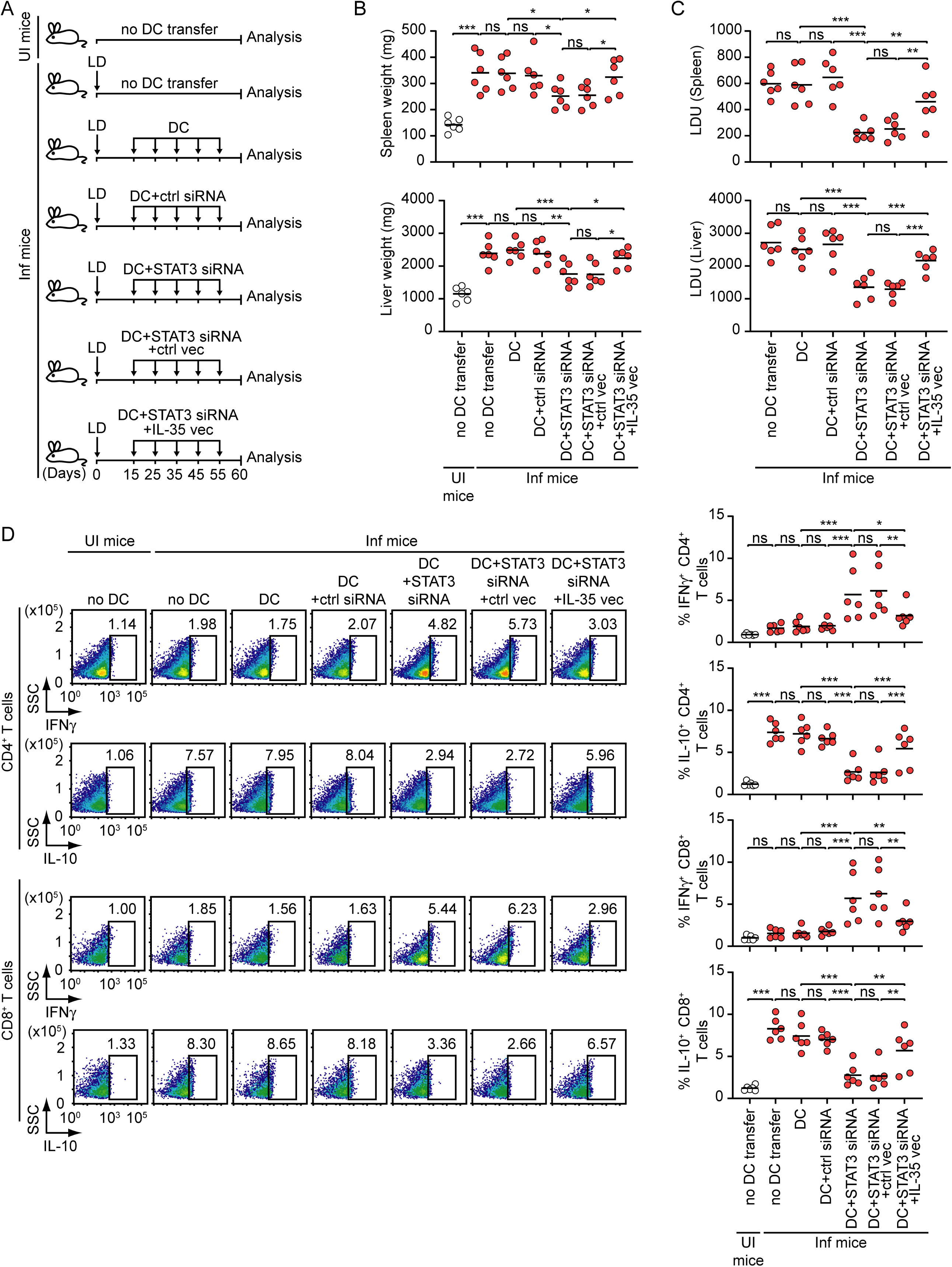
The STAT3-IL-35 axis in DCs drives immunosuppression and disease pathogenesis during LD infection. (**A**) Protocol for adoptive DC transfer: BALB/c BMDCs (1 x 10^6^) were transfected with STAT3 siRNA [either alone or in combination with control vector (Ctrl vec) or IL-35-expressing vector (IL-35 vec); IL-35 expression in these BMDCs is shown in Fig. EV8] or with control siRNA. These cells were then administered intravenously to LD-infected BALB/c mice on the indicated days after LD infection. In some groups, DC transfer was not performed in uninfected or LD-infected mice. At day 60 postinfection, spleen and livers were subjected to the analyses as described in (B to D). (**B** and **C)** Spleen and liver weights (B) and parasite burdens (C). Data are combined from two experiments (*n* = 3 mice per group in each experiment). (**D)** Frequencies of IFNγ- and IL-10-expressing CD4^+^ and CD8^+^ T cells in the spleen were assessed by flow cytometry (representative of *n* = 6; left). Right: pooled results from two experiments (*n* = 3 mice per group per experiment). The gating strategy was same as in Fig. EV6A. In (B), (C), and (D, right panel), each symbol represents the data of an individual mouse, and horizontal bars denote mean values. \**P* < 0.05, \*\**P* < 0.01, \*\*\**P* < 0.001; ns, not significant.

**Figure 6.**
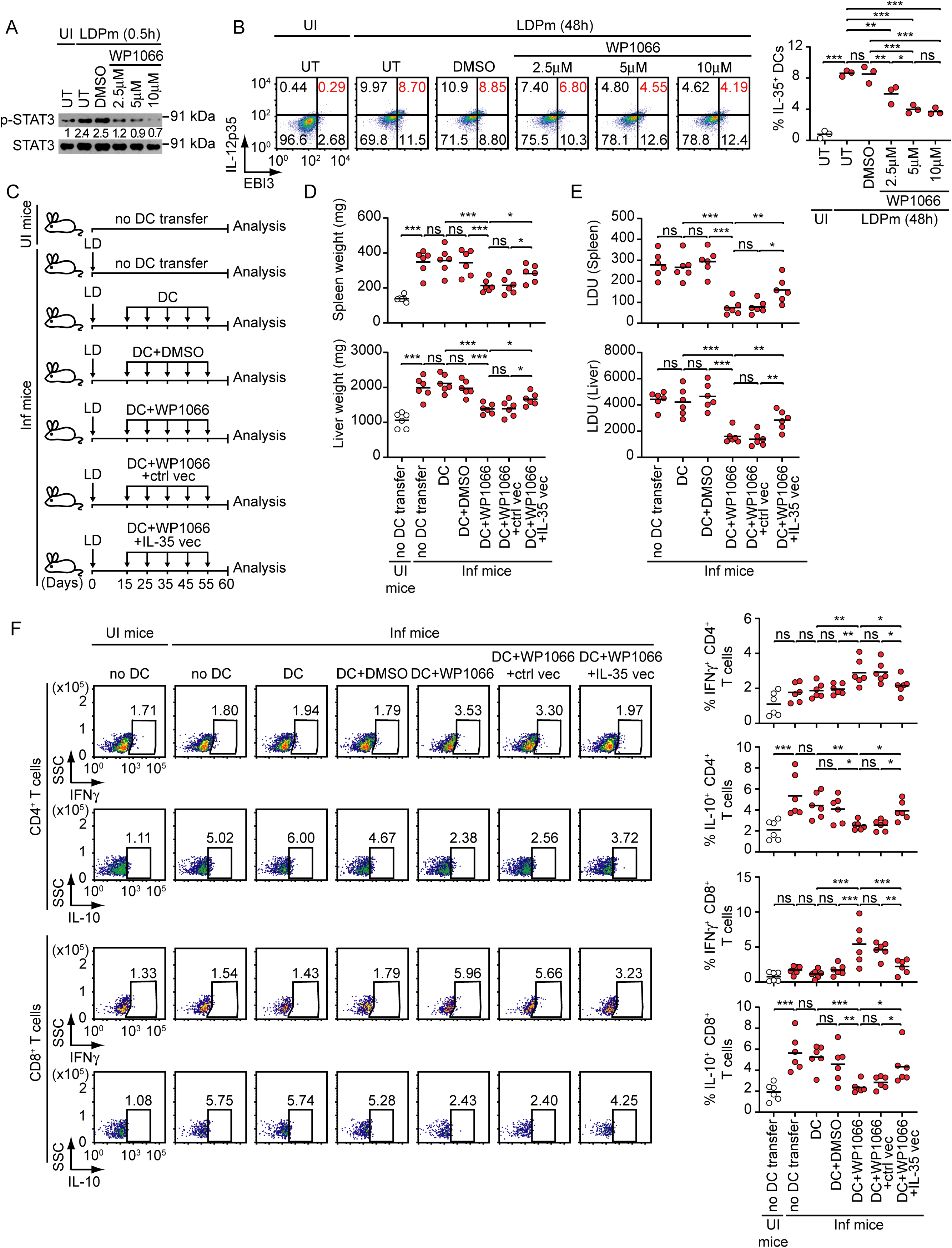
Inhibition of DC IL-35 production by WP1066 augments anti-leishmanial immunity. (**A**) Expression of phosphorylated and total STAT3 (loading control) in uninfected and LDPm-infected (0.5 h) BMDCs that were left untreated (UT) or treated with DMSO (0.1%) or WP1066 (concentrations indicated above the lanes) for 1 h was determined by immunoblotting (representative of *n* = 3 experiments). Relative densitometry values (normalized to total STAT3) are shown below the lanes. Combined densitometry data from *n* = 3 experiments are presented in Fig. EV9A. **B** The frequency of IL-35-expressing BMDCs (uninfected and LDPm-infected for 48 h) treated as in (A) was assessed by flow cytometry (representative of *n* = 3 experiments; left). Right: combined data from *n* = 3 experiments. Cytotoxicity of DMSO and WP1066 is shown in Fig. EV9B. (**C**) Procedure for transfer of WP1066-treated DCs: BALB/c BMDCs (1 x 10^6^), transfected (or not) with control vector (Ctrl vec) or IL-35-expressing vector (IL-35 vec), were treated for 1 h with DMSO (0.1%) or WP1066 (5 μM) or left untreated, as indicated (see Fig. EV9C for IL-35 expression in these BMDCs). BMDCs were then transferred to LD-infected BALB/c mice on different days (15-55 days) after infection. At day 60 postinfection, spleens and livers were collected for the analyses shown in (D to F). (**D** and **E**) Liver and spleen weights (D) and parasite burdens (E) in the mice described above. Data from two experiments are summarized (*n* = 3 mice per group in each experiment). (**F**) Flow cytometric analysis of IFNγ- and IL-10-expressing CD4^+^ and CD8^+^ T cells in the spleen of the mice described in (C). Left: representative plots out of *n* = 6. Right: summary of two experiments (*n* = 3 mice per group in each experiment). The gating strategy was the same as that described in Fig. EV6A. In (B, right panel), each symbol represents the data of an independent experiment; in (D), (E), and (F, right panel) each symbol represents the data of an individual mouse. Horizontal bars in these panels indicate the mean values. \**P* < 0.05, \*\**P* < 0.01, \*\*\**P* < 0.001; ns, not significant.

### Reducing DC IL-35 production by targeting STAT3 promotes parasite clearance *in vivo*

We next investigated whether lowering DC-derived IL-35 production by suppressing STAT3 could limit disease pathogenesis. To test this, we adoptively transferred either unsilenced (control siRNA-transfected) or STAT3-silenced BMDCs into LD-infected mice at different days postinfection (Fig. 5A). At day 60 postinfection, we isolated the spleens and livers of these mice and found that adoptive transfer of STAT3-silenced BMDCs, but not unsilenced BMDCs, significantly reduced liver and spleen weights as well as parasite burden in LD-infected mice (Fig. 5, B and C). To further verify whether these effects of STAT3-silenced BMDCs were due to decreased IL-35 production, we overexpressed IL-35 in these cells (Fig. 5A and Fig. EV8). Overexpression of IL-35 considerably attenuated the capacity of STAT3-silenced BMDCs to reduce liver and spleen weights and parasite load in LD-infected mice (Fig. 5, B and C). Consistent with these findings, transfer of STAT3-silenced BMDCs increased the frequency of IFNγ-expressing and decreased the frequency of IL-10-expressing CD4^+^ and CD8^+^ T cells in the spleens of LD-infected mice, whereas IL-35 overexpression reversed these immunological outcomes (Fig. 5D). These results suggest that targeting STAT3 to inhibit DC-derived IL-35 production promotes parasite clearance *in vivo* and restrains disease pathogenesis.

We then investigated whether WP1066, an FDA-approved orphan drug and STAT3 inhibitor (Wang *et al*., 2022) (https://www.moleculin.com/technology/wp1066/), can suppress LD-induced IL-35 production in DCs and limit subsequent disease pathogenesis. Accordingly, we first examined the effect of WP1066 treatment on LD-induced STAT3 activation and IL-35 production in BMDCs. While LDPm readily induced STAT3 activation and IL-35 expression in untreated and DMSO (0.1%, control)-treated BMDCs, these effects were markedly reduced in BMDCs treated with 2.5-10 μM WP1066 (Fig. 6, A and B, and Fig. EV9A). Notably, WP1066 showed almost no toxicity at all tested concentrations (2.5-10 μM), as evidenced by the negligible frequency of dead cells (Fig. EV9B). For subsequent experiments, however, we used 5 μM WP1066, the lowest concentration that maximally suppressed IL-35 production while causing insignificant cell death compared to untreated BMDCs (Fig. 6B and Fig. EV9B). Collectively, the above results demonstrate that WP1066 suppresses LD-induced STAT3 activation and IL-35 production by DCs. We then assessed whether WP1066-mediated inhibition of DC IL-35 production could limit disease progression. To test this, we adoptively transferred untreated, DMSO-treated, or WP1066-treated BMDCs (with or without IL-35 overexpression) into LD-infected syngeneic mice at the indicated days after LD infection (Fig. 6C). At day 60 postinfection, we found that transfer of WP1066-treated BMDCs, but not untreated or DMSO-treated BMDCs, reduced liver and spleen weights and parasite burden (Fig. 6, D and E). Moreover, transfer of WP1066-treated BMDCs decreased the frequency of IL-10-expressing CD4^+^ and CD8^+^ T cells and increased the frequency of IFNγ-expressing CD4^+^ and CD8^+^ T cells in the spleens of LD-infected mice (Fig. 6F). Strikingly, IL-35 overexpression attenuated the ability of WP1066-treated BMDCs to mediate these effects (Fig. 6, D to F, and Fig. EV9C). These data confirm that WP1066 effectively inhibits STAT3-mediated IL-35 production by DCs and prevents disease exacerbation. Furthermore, together with our other findings that oral administration of WP1066 significantly reduced liver and spleen weights and parasite burden in LD-infected mice (Fig. 7), these results highlight WP1066 as a potential new therapeutic agent against VL.

**Figure 7.**
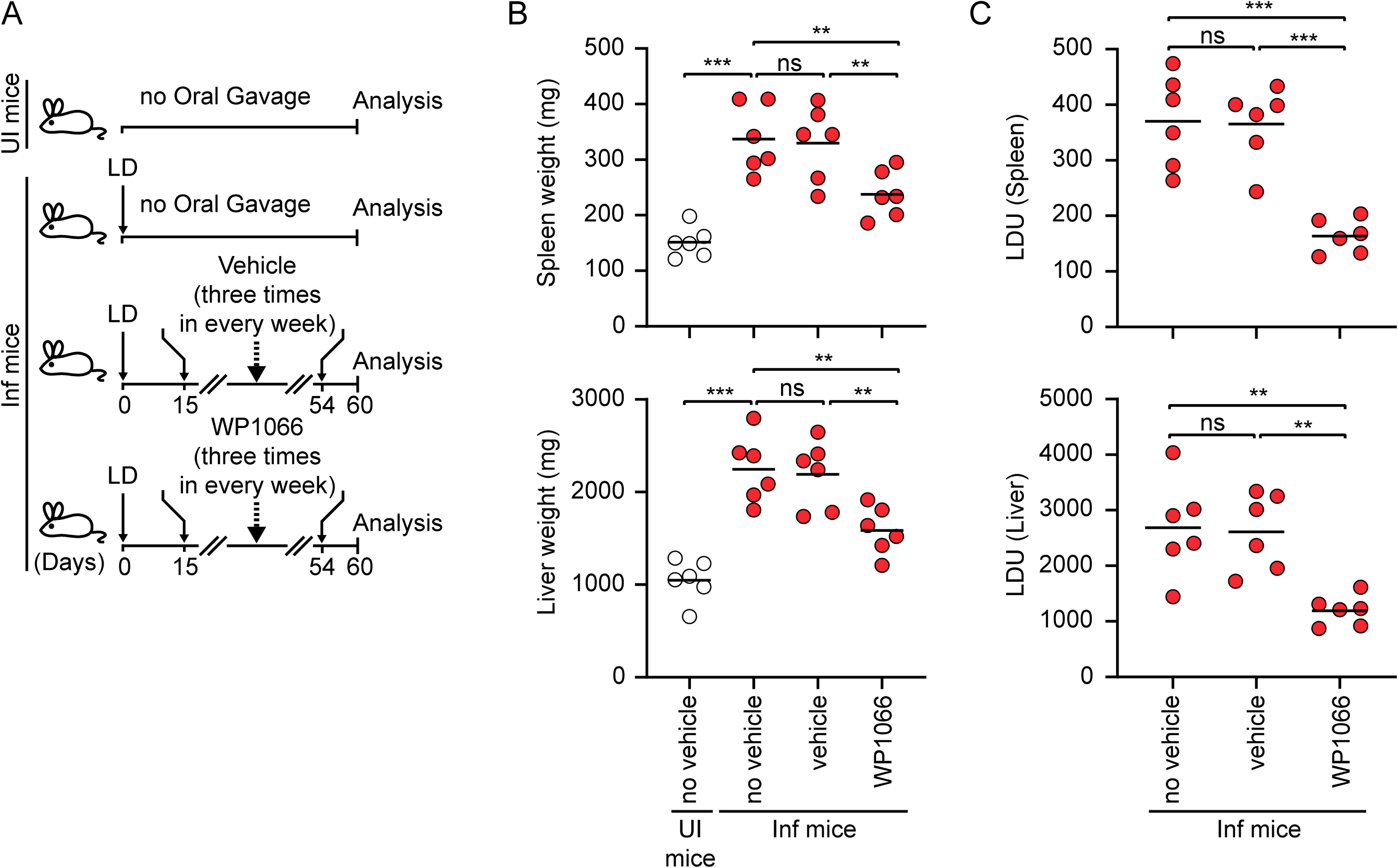
Oral administration of WP1066 reduces LD infection in mice. (**A**) WP1066 administration protocol: WP1066 (20 mg/kg body weight) or the control vehicle (100 μl PEG 300/DMSO) was administered orally to LD-infected BALB/c mice three times weekly (i.e. on alternate days) from day 15 to day 54 postinfection. On day 60 postinfection, spleens and livers were collected to assess parasite burden. Uninfected mice, LD-infected mice without oral gavage, and LD-infected mice receiving only the control vehicle served as controls. (**B** and **C)** Spleen and liver weights (B) and parasite burdens (C) in the mice describes above. Data from two experiments (*n* = 3 mice per group in each experiment) were combined. Each symbol in (B) and (**C**) represents the data from a single mouse, horizontal bars indicate mean values.\*\**P* < 0.01, \*\*\**P* < 0.001; ns, not significant

## DISCUSSION

The current study reveals a pivotal role for IL-35 in driving immunosuppression and the pathogenesis of LD infection. It also identifies DCs as the initial source of IL-35, defines the receptor and the signaling pathway that mediate IL-35 induction in these cells, and uncovers a promising therapeutic compound that suppresses both IL-35 production and LD infection. Five major findings emerge from this work.

First, LD stimulates IL-35 production in DCs. Previous studies, based on the expression of IL-12p35 and EBI3, had claimed that IL-35 is produced by tolerogenic (dexamethasone-treated) and mature (LPS-treated) DCs (Dixon *et al*., 2015; Liu *et al*, 2018). However, these studies did not investigate whether IL-12p35 and EBI3 are co-expressed in the same DC and whether these two subunits physically interact to form the IL-35 heterodimer. Another study reported that increased EBI3 expression in CD4^+^ T cells promotes immunosuppression and contributes to LD pathogenesis (Asad *et al*., 2019); however, the role of the complete IL-35 heterodimer was not investigated. Notably, IL-12p35 and EBI3 alone can also exert immunosuppressive effects (Hildenbrand *et al*., 2023). Therefore, the production of IL-35 by DCs and its role in VL have remained undefined. Moreover, it is not known whether DCs can produce IL-35 in response to a pathogenic challenge. Our results, supported by flow cytometry, confocal microscopy, ELISA, c-FRET, and co-immunoprecipitation analyses, demonstrate that DCs can indeed produce the IL-35 heterodimer, albeit in response to LD infection. These findings identify DCs as a novel source of IL-35 in the context of infectious disease.

Second, DCs act as the initial producers of IL-35 during LD infection. The IL-35-expressing cells in the spleen began to increase as early as 22 days after LD infection. At this time point, the frequency of IL-35-producing DCs was substantially higher than that of IL-35-producing T cells or other splenocytes. Moreover, IL-12p35 and EBI3 levels were higher in IL-35-producing DCs compared with IL-35-producing T cells or other splenic cells. These observations establish DCs as the primary early source of IL-35 during LD infection. Notably, DC-derived IL-35 can, in turn, induce IL-35 expression in T cells. For instance, culture supernatants from LD-infected DCs increased the frequency of IL-35-expressing T cells in an IL-35-dependent manner. Likewise, adoptive transfer of DCs transfected with IL-35-neutralizing antibody reduced the frequency of IL-35-expressing T cells in LD-infected mice, providing direct *in vivo* evidence that DC-derived IL-35 drives the expansion of IL-35-producing T cells during LD infection. Collectively, these results reveal an IL-35-dependent cross-talk between DCs and T cells, through which DCs may initiate and shape the immunosuppressive environment during LD infection. Furthermore, our findings demonstrating the ability of DC-produced IL-35 to induce IL-35 expression in T cells provide the first evidence of this regulatory interaction not only in the context of LD infection but also in infectious diseases more broadly.

Third, IL-35 expressed during LD infection promotes disease pathogenesis and DCs, as the initial IL-35 producers, play a crucial role in this process. These conclusions are supported by our observations that neutralization of IL-35 in LD-infected mice or in adoptively transferred DCs (via *in vivo* administration or transfection of anti-IL-35 antibody, respectively) reduced parasite burden in the liver and spleen and enhanced the anti-leishmanial immune response. Notably, the IL-12p35 or EBI3 knockout mice were not suitable for these studies. This is because IL-12p35 and EBI3 alone can act as immunosuppressors (Hildenbrand *et al*., 2023). Accordingly, in such knockout models, it would remain unclear whether the observed effects were due to the absence of IL-12p35 or EBI3 individually, or to the absence of the complete heterodimeric IL-35 complex. There is also evidence that IL-12p35 deficiency impairs both IL-12 and IL-35 expression (Huang *et al*, 2019), and EBI3 deficiency prevents both IL-35 and IL-27 expression (Liu *et al*, 2015). These reports suggest that deficiency of IL-12p35 or EBI3 affects not only IL-35 expression, but also the expression of other related cytokines such as IL-12 and IL-27. Therefore, neutralization of IL-35 with an anti-IL-35 antibody provided only reliable approach to assess the contribution of IL-35 to immune regulation during LD infection. Importantly, the anti-IL-35 antibody (clone V1.4C4.22) used here selectively neutralizes IL-35 and not IL-27, IL-12 (Collison *et al*., 2010) or EBI3 (Fig. EV10). Together, our findings that IL-35, particularly the one produced by DCs during LD infection, inhibits the anti-leishmanial immune response and promotes disease pathogenesis reveal a previously unidentified immunosuppressive mechanism associated with LD infection. We further demonstrate that IL-35 released by DCs during LD infection suppresses anti-leishmanial immunity by inhibiting DC activation and maturation (via suppression of NF-κB signaling) and by dampening T cell responses. Overall, these results elucidate the mechanism by which DC-derived IL-35 suppresses anti-leishmanial immune response.

Fourth, our study shows that TIM-3-STAT3 signaling plays a pivotal role in LD-induced IL-35 production in DCs. When TIM-3 was blocked with an anti-TIM-3 antibody or STAT3 expression was suppressed, IL-35 expression was greatly reduced in DCs despite LD infection. Subsequent analysis showed that STAT3 mediates IL-35 induction by directly binding to the *IL12A* and *EBI3* promoters. Our findings, however, do not rule out the possibility of a STAT3-independent mechanism contributing to IL-35 expression in DCs. Currently, the role of TIM-3 and STAT3 in regulating IL-35 expression is not known, although other STAT family members, such as STAT1 and STAT4, have been reported to drive IL-35 expression in conventional T cells (Collison *et al*., 2012). Thus, our results reveal a new role for TIM-3 and STAT3 in promoting IL-35 expression. Moreover, our results indicate that STAT3-mediated IL-35 production by DCs contributes to immunosuppression and disease pathogenesis during LD infections. For example, adoptive transfer of STAT3-silenced DCs markedly reduced the immunosuppressive type-2 T cell response, hepatosplenomegaly, and parasite burden in LD-infected mice; however, these effects were attenuated when IL-35 was overexpressed in these cells. Given that STAT3 is essential for enhanced IL-35 production and consequent disease pathogenesis, we further explored a STAT3-targeted therapeutic approach to limit IL-35 expression, enhance the anti-leishmanial immune response and reduce LD infection.

Fifth, we show that the FDA-approved orphan drug and STAT3 inhibitor WP1066 reduces IL-35 production in DCs during LD infection, thereby enhancing the host-protective type-1 T cell response and lowering the parasite burden in LD-infected mice. In fact, oral administration of WP1066 to LD-infected mice markedly reduced the parasite load. Although previous studies have reported the anti-cancer properties of WP1066 (Hussain *et al*, 2007; Iwamaru *et al*, 2007; Tsujita *et al*, 2017), its potential role in combating pathogenic infections has remained unexplored. Our study provides the first evidence that WP1066 limits LD infection, highlighting its strong potential for repurposing against VL and paving the way for future research into its therapeutic potential against other infectious diseases. Furthermore, the identification of WP1066 as a suppressor of IL-35 production is particularly significant, as no IL-35 suppressor is currently known.

Overall, this study identifies a novel role for DCs as the initial producers of IL-35 during LD infection and establishes DC-derived IL-35 as a key driver of immunosuppression and disease pathogenesis. LD induces IL-35 expression in DCs by activating STAT3 through the TIM-3 receptor. In this context, we have previously shown that Bruton’s tyrosine kinase (Btk), a member of the Tec kinase family, acts as a signaling link between TIM-3 and STAT3 (Mishra *et al*., 2023). Activated STAT3 then binds to the *IL12A* and *EBI3* promoters, thereby promoting IL-35 expression in DCs. The IL-35 produced by DCs subsequently suppresses DC and T cell activation, enhances IL-35 expression in T cells, inhibits the anti-leishmanial immune response and ultimately promotes disease pathogenesis. Finally, we demonstrated that the STAT3 inhibitor WP1066 suppresses IL-35 production and significantly reduces LD infection *in vivo* (Fig. 8). Together, these findings delineate a unique immunosuppressive mechanism that contributes to VL pathogenesis and highlight WP1066 as a promising therapeutic candidate. Moreover, our study provides a strong rationale for future clinical evaluation of WP1066 as an anti-*Leishmania* agent and opens avenues to explore the broader role of DC-derived IL-35 in immunosuppression and the pathogenesis of other infectious diseases.

**Figure 8.**
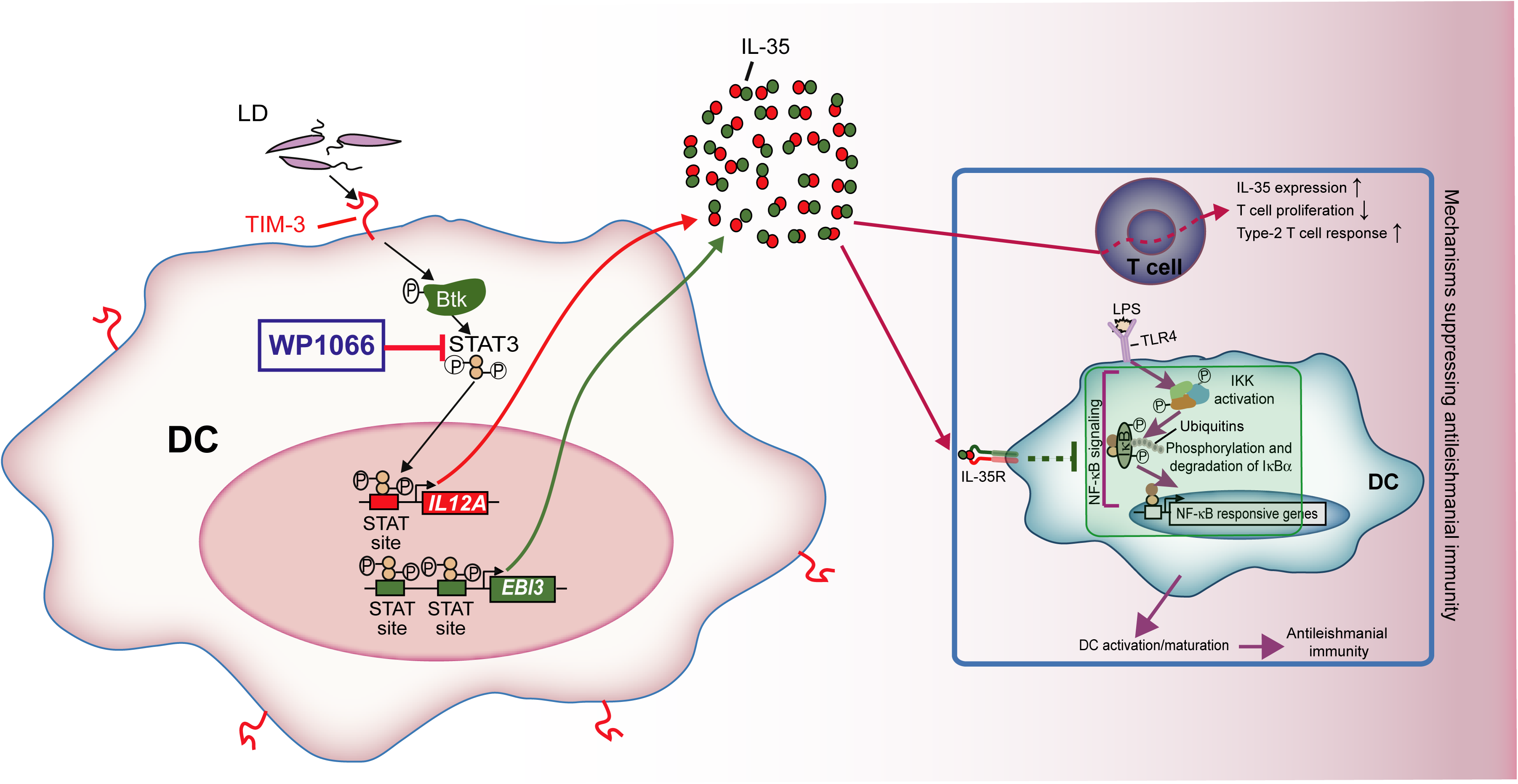
A model illustrating how LD induces IL-35 production in DCs and thereby suppresses anti-leishmanial immunity. LD activates STAT3 in DCs via the TIM-3 receptor and its downstream signaling mediator Btk (as shown in our previous report (Mishra *et al*., 2023)). Activated STAT3 then directly promotes IL-35 production in DCs by binding to the *IL12A* and *EBI3* promoters (*IL12A* and *EBI3* encode the IL-35 subunits IL-12p35 and EBI3, respectively). DC-derived IL-35, in turn, inhibits the activation and maturation of bystander DCs by suppressing the NF-κB signaling pathway (inner green shaded box), promotes IL-35 expression in T cells, reduces T cell proliferation, and drives pathogenic type-2 T cell responses. Collectively, these events (summarized in the blue-outlined box) impair anti-leishmanial immunity and exacerbate disease pathogenesis. Notably, pharmacological blockade of STAT3 activation by WP1066 reduces IL-35 production by DCs, suppresses disease-promoting type-2 T cell responses, enhances host-protective type-1 T cell responses, and ultimately lowers parasite burden *in vivo*.

## MATERIALS AND METHODS

### Study Approval

The Institutional Animal Ethics Committee at the CSIR-Institute of Microbial Technology approved all animal experiments (IAEC/21/16, IAEC/23/15), which were conducted as per the National Regulatory Guidelines of the CPCSEA (Committee for the Purpose of Control and Supervision of Experiments on Animals). The Institutional Biosafety Committee has approved the studies involving LD (CSIR-IMTECH/IBSC/2021/June 06, CSIR-IMTECH/IBSC/2024/July 13).

### Reagents

Immunoblot analyses were performed with the following antibodies and reagents: anti-IKKα (sc-7182), anti-IKKβ (sc-34673), anti-IκBα (sc-371), anti-β-actin (sc-47778), HRP-anti-rat IgG (sc-2032) and HRP-anti-goat IgG (sc-2354; all from Santa Cruz Biotechnology, TX, USA); anti-phospho-IKK (Ser180/181, SAB4301429; Sigma-Aldrich, India); anti-phospho-STAT3 (Tyr705, 9131) and anti-STAT3 (9139; both from Cell Signaling Technologies, MA, USA); biotin-anti-EBI3 (210-306-B66), anti-IL-12p35 (C18.2; 14-7122-85) and HRP-Streptavidin (N100; all from Thermo Fisher Scientific, MA, USA); and HRP-anti-mouse IgG (HAF007) and HRP-anti-rabbit IgG (HAF008 both from R&D Systems, MN, USA). Immunoprecipitation was performed with the following antibodies: anti-EBI3 (MABF848; Sigma-Aldrich); and mouse IgG2b (02-6300), anti-IL-12p35 (PA5-79460) and rabbit IgG (02-6102; all from Thermo Fisher Scientific). For ChIP assays, the following antibodies and reagents were used: anti-STAT3 (4904) and rabbit IgG (2729; both from Cell Signaling Technologies); and protein A/G PLUS-Agarose beads (sc-2003; Santa Cruz Biotechnology). The following antibodies were used for flow cytometry: PE-anti-mouse EBI3 (IC18341P) and APC-anti-human/mouse IL-12p35 (IC2191A) (both from R&D Systems); eFluor 660-anti-mouse IL-12p35 (50-7352-82), PerCP-anti-mouse/human IL-12p35 (MA5-23622) and Alexa Fluor 594-anti-mouse IgG (A-11020; all from Thermo Fisher Scientific); PE-anti-human EBI3 (360903), FITC-anti-mouse CD40 (124608), FITC-anti-mouse CD86 (105006), FITC-anti-mouse CD80 (104706), FITC-anti-mouse CD11c (117306), PE-anti-mouse CD8α (100708), FITC-anti-mouse CD8α (100706), PE/Cyanine7-anti-mouse CD3 (100220), PerCP/Cyanine5.5-anti-mouse CD4 (100434), Alexa Fluor 647-anti-mouse IDO1 (654003), APC-anti-mouse IL-10 (505010), FITC-anti-mouse IFNγ (505806) and isotype control antibodies such as FITC-rat IgG2a,κ (400505) and FITC-armenian hamster IgG (400905; all from Biolegend, CA, USA). Neutralization studies were performed using neutralizing anti-IL-35 (clone V1.4C4.22, MABF848; Sigma-Aldrich) and the corresponding isotype control mouse IgG2b (BE0086; Bio X Cell, NH, USA). The TIM-3 blocking antibody (clone RMT3-23, BE0115) (Chiba *et al*., 2012) and the isotype control rat IgG2a,κ (BE0089) were obtained from Bio X Cell. The following antibodies were used for confocal microscopy: PE-anti-mouse EBI3 (IC18341P) and APC-anti-human/mouse IL-12p35 (IC2191A; both from R&D Systems). For c-FRET analysis, the following antibodies and reagents were used: biotin-anti-EBI3 (210-306-B66), Alexa Fluor 568-Streptavidin (S11226), anti-IL-12p35 (C18.2; 14-7122-85) and FITC-anti-rat IgG (31629; all from Thermo Fisher Scientific). Anti-mouse CD3ε (100340) and anti-mouse CD28 (102116) antibodies used for T cell proliferation assay were obtained from BioLegend. The following recombinant proteins were used in this study: Mouse granulocyte-macrophage colony-stimulating factor (GM-CSF) and mouse IL-4 were obtained from Peprotech Asia (Rehovot, Israel); human GM-CSF and human IL-4 were purchased from Miltenyi Biotec (Germany); IL-35 (mouse)-IgG1 Fc (human) (CHI-MF-11135-C025) and human IgG1 Fc (control; CHI-HF-210IG1-C100) were procured from Chimerigen/Biomol (Hamburg, Germany); and mouse EBI3 (CYT-621) was purchased from Prospec (Rehovot, Israel). The ON-TARGETplus SMARTpool siRNAs targeting mouse STAT3 and non-targeting control pool siRNAs were procured from Dharmacon (CO, USA). Anti-mouse CD3ε magnetic microbeads (130-094-973) were obtained from Miltenyi Biotec. The Dynabeads M-280 Streptavidin for DNA pull-down assays was from Thermo Fisher Scientific. The eFluor 670 dye (65-0840-85) and Intracellular Fixation & Permeabilization Buffer kit with Brefeldin A (88-8823-88) were also purchased from Thermo Fisher Scientific. WP1066 (HY-15312) was obtained from MedChemExpress (NJ, USA), and LPS (*Escherichia coli* O111:B4) and other reagents from Sigma-Aldrich.

### Animals and Parasites

BALB/c mice and golden hamsters (*Mesocricetus auratus*) were housed and bred in a pathogen-free environment at IMTECH Centre of Animal Resources and Experimentation (iCARE). The LD parasite [strain AG83 (MHOM/IN/83/AG83); American Type Culture Collection (ATCC PRA-413)] was regularly maintained in golden hamsters (Chakraborty *et al*, 2005). Amastigotes were obtained from infected hamster spleens as previously described (Hart *et al*, 1981) and used in some experiments. Alternatively, after isolation, amastigotes were transformed into promastigotes (Chakraborty *et al*., 2005), which were then cultured and used in experiments.

### DC preparation, LD infection and treatments

BMDCs were generated from bone marrow cells of male or female BALB/c mice (8–12 weeks old) as described (Wallet *et al*, 2008). BMDCs (5 x 10^6^/well unless otherwise indicated) were then infected *in vitro* with either LDAm or stationary phase LDPm at a DC to parasite ratio of 1:10 for the indicated times in RPMI 1640 complete medium (10% FBS, penicillin/streptomycin, Na-pyruvate, L-glutamine and 2-mercaptoethanol). Subsequently, the BMDCs were washed and in certain cases stimulated with LPS (500 ng/ml). In some studies, BMDCs were treated with different concentrations of the STAT3 inhibitor WP1066 (2.5-10 μM) or DMSO (0.1%; control) for 1 h prior to LDPm infection. The cytotoxic effect of a wide range of WP1066 concentrations (2.5-100 μM) and 1% DMSO on BMDCs was assessed by staining with propidium iodide (PI) and assessing the frequency of dead cells (PI^+^ BMDCs) by flow cytometry as described (Qin *et al*, 2022) (Fig. EV9B). For TIM-3 blocking experiments, BMDCs were treated with 10 μg/ml anti-TIM-3 antibody (clone RMT3-23) or an isotype control antibody before incubation with LDPm as described (Akhtar *et al*., 2022). Notably, some experiments were performed using HuMoDCs (CC-2701, Lonza, Basel, Switzerland) (Akhtar *et al*., 2020) instead of murine DCs.

### Analysis of intracellular IL-35 expression

BMDCs (1 x 10^6^ cells/well) were infected *in vitro* with LDPm or LDAm for the indicated time periods. Brefeldin A (10 μg/ml) was then added 3 h before harvesting the cells. Subsequently, intracellular immunostaining was carried out using the Intracellular Fixation and Permeabilization Kit (eBioscience) and the antibodies PE-anti-EBI3 and PerCP- or APC-anti-IL-12p35. The frequency of IL-35-expressing BMDCs (IL-12p35^+^EBI3^+^ BMDCs) was determined by flow cytometry. An assessment of IL-35 expression in splenic DCs, T cells and other splenocytes (i.e., non-DC, non-T cell populations) from uninfected and LD-infected mice was also performed. For this purpose, splenocytes isolated from LD-infected mice (at the indicated postinfection time points) and uninfected mice were stained with FITC-anti-CD11c (for sDCs) or PE/Cyanine7-anti-CD3 (for T cells). Intracellular immunostaining of EBI3 and IL-12p35 was then done as described above. The frequency of IL-35-expressing (IL-12p35^+^EBI3^+^) sDCs (CD11c^+^ cells), T cells (CD3^+^ cells) and other splenocytes (CD11c^-^CD3^-^ cells) and the levels of IL-35 subunits (IL-12p35 and EBI3) expressed in these cells were determined by flow cytometry. In some experiments, intracellular immunostaining of IDO with Alexa Fluor 647-anti-mouse IDO1 antibody was performed to determine the expression of IDO in IL-35-expressing sDCs.

### Confocal microscopy and c-FRET analysis

BMDCs (2.5 × 10^5^) were seeded on 12-mm coverslips, allowed to adhere for 6 h and then infected with LDPm for 48 h. Cells were stained with the primary antibodies anti-IL-12p35 and biotin-anti-EBI3 (each at a dilution of 1:500) for 1 h at room temperature as previously described (Akhtar *et al*., 2020). The cells were then washed and incubated with the secondary antibody or reagent FITC-anti-rat IgG and Alexa Fluor 568-Streptavidin (both at a dilution of 1:1000) for 1 h at room temperature. The cell nuclei were stained with Hoechst (1 mg/ml; Thermo Fisher Scientific). Images were captured using a Nikon A1R laser scanning confocal microscope and processed with Fiji and Adobe Photoshop. Colocalization efficiency was measured by Pearson’s Coefficient and Manders’ Coefficient as described (Dunn *et al*, 2011). To perform the acceptor photobleaching assay, a 25-second pulse of high-intensity laser (561 nm) was applied to bleach the Alexa Fluor 568 signal (acceptor) within the cells. A series of fluorescence intensity measurements for FITC (donor) were performed before and after bleaching. The percentage of FRET efficiency (E) was calculated using the following equation: 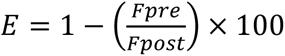; where F_pre_ and F_post_ represent the fluorescence intensity of the donor before and after photobleaching of the acceptor, respectively.

### Treatment of DCs and T cells with LD-infected DC supernatants

In this study, the culture supernatant of BMDCs (5 x 10^6^ cells in 3 ml RPMI complete medium per well) infected with LDPm for 48 h was used to analyze the effect of IL-35, if present in the supernatant, on DCs and T cells. The culture supernatant of uninfected BMDCs served as control. To determine the effect on DCs, BMDCs from uninfected mice were cultured for 24 h in a 6-well ultra-low attachment plate (Sigma-Aldrich) in the presence or absence of the above supernatants (50% of total medium) and 10 μg/ml of neutralizing anti-IL-35 or isotype control antibody. Subsequently, BMDCs were activated with LPS for 24 h. The expression of costimulatory molecules (CD40, CD80 and CD86) and the secretion of cytokines (IL-12 and TNFα) by BMDCs were analyzed by flow cytometry and ELISA, respectively. In addition, the activation of NF-κB signaling pathway in BMDCs was examined via electrophoretic mobility shift assay (EMSA) and immunoblot analysis.

To determine the effect on T cells, T cells were purified from spleens of uninfected mice using anti-CD3 magnetic microbeads. Cells (1 x 10^6^ cells/well) were then cultured in a 24-well plate (pre-coated overnight with 2 µg/ml anti-CD3 and anti-CD28 antibodies) in the presence or absence of uninfected or LD-infected BMDC culture supernatants together with 10 μg/ml neutralizing anti-IL-35 or isotype control antibody. After 72 h, cells were harvested and stained with PerCP/Cyanine5.5-anti-CD4 or FITC-anti-CD8 antibodies. The expression of IL-35 in CD4^+^ and CD8^+^ T cells was analyzed by flow cytometry as mentioned above. In some experiments, T cells purified from uninfected mice were labeled with eFluor 670 dye (5 μM) and cultured as described above. Alternatively, eFluor-labeled T cells were cultured in the presence of recombinant mouse EBI3, mouse IL-35-human IgG1 Fc or human IgG1 Fc control (in all cases 10 μg/ml; instead of BMDC supernatants) and 10 μg/ml neutralizing anti-IL-35 antibody or mouse IgG2b (isotype control). After 72 h, the proliferation of T cells was assessed by flow cytometry based on the dilution of eFluor 670 dye fluorescence intensity.

### Quantitative PCR (qPCR) and RT-qPCR analyses

Isolation of genomic DNA (from the splenic suspension of infected mice) and total RNA (from DCs) was performed using the DNeasy Blood and Tissue kit (Qiagen, Germany) and the RNA extraction kit (Promega, WI, USA), respectively. The qPCR and RT-qPCR analyses were performed using the GoTaq qPCR and RT-qPCR systems (Promega) and primers listed in table EV1 following the manufacturer’s guidelines. PCR reactions were conducted using the qTOWER³ real-time PCR thermal cycler (Analytik Jena, India) or the CFX96 Touch Real-Time PCR Detection System (Bio-Rad, CA, USA). The mRNA expression of target genes was quantified by RT-qPCR as described (Gujar *et al*, 2016).

### ChIP

ChIP was performed with anti-STAT3 antibody or rabbit IgG using the ChIP-IT kit (Active Motif, CA, USA). The immunoprecipitated DNA fragments were then analyzed by qPCR to determine STAT3 recruitment to the mouse *EBI3* and *IL12A* promoters (see table EV1 for primer details). Results were normalized to those obtained with rabbit IgG (nonspecific background) and presented as fold enrichment of STAT3 relative to uninfected DCs.

### EMSA, DNA pull-down assay, immunoprecipitation and immunoblot analysis

Nuclear extracts were made from DCs as mentioned previously (Beg *et al*, 1993). EMSA was performed using a ^32^P-labeled H2K (MHC-I) promoter-specific DNA probe containing NF-κB-binding sites, 5′-CAGGGCTGGGGATTCCCCATCTCCACAGTTTCACTTC-3′, or an OCT-1 DNA probe (control), 5′-TGTCGAATGCAAATCACTAGAA-3′ (Akhtar *et al*., 2022). Bands were visualized using a phosphorimager (PharosFX Molecular Imager; Bio-Rad).

DNA pull-down assay of DC nuclear extracts was performed using various biotinylated wild-type or mutant oligonucleotides (table EV1) and streptavidin-coupled Dynabeads as described (Chan *et al*, 2009). Immunoblot analysis of the pull-down proteins was then performed as previously illustrated (Bhattacharyya *et al*, 2004).

Immunoprecipitation was performed as described (Akhtar *et al*., 2022). Briefly, uninfected or LDPm-infected DCs were treated with the protein cross-linker dimethyl 3,3-dithiopropionimidate dihydrochloride (2 mg/ml, Sigma-Aldrich) and then lysed as described (Akhtar *et al*., 2022). The DC lysates were then immunoprecipitated with anti-EBI3, anti-IL-12p35, or the corresponding control antibodies (mouse IgG2b or rabbit IgG) and analyzed by immunoblotting.

### Densitometry

Scion Image software (Scion Corporation, Maryland, USA) was used for densitometric quantification.

### Reporter assay

The pGL3-firefly luciferase reporter constructs (Promega, WI, USA) with wild-type *IL12A* promoter region [-391/-1 region; *IL12A* (Wt)-pro] or wild-type *EBI3* promoter region [-412/-1 region; *EBI3* (Wt)-pro] and their various mutant versions for the STAT-binding sites [*IL12A* (Mut)-pro, *EBI3* (Mut1)-pro and *EBI3* (Mut2)-pro] were synthesized by Biotech Desk Pvt. Ltd (India). The *EBI3* (Mut1)-pro and *EBI3* (Mut2)-pro constructs contained mutations at two different STAT binding sites. Each of these reporter constructs (400 ng) was individually transfected into HEK293T cells (2.5 x 10^5^ cells/well), along with 400 ng of pBabe-mouse STAT3 or empty pBabe vector and 200 ng of pRL-CMV plasmid [expresses *Renilla* luciferase (internal control; Promega)] using FreeStyle MAX reagent (Thermo Fisher Scientific). After 24 h, luciferase activity was measured using the Dual-Luciferase Reporter Assay Kit (Promega).

### DC transfection

BMDCs (2.5 x 10^5^) were transfected with siRNAs using Lipofectamine RNAiMAX reagent (Thermo Fisher Scientific) as previously described (Akhtar *et al*., 2022). Transfection of DNA constructs such as mouse IL-35-expressing vector (pIGneo-IL-35-GFP; 2 μg) or empty vector (pIGneo-GFP; 2 μg) into BMDCs (1 x 10^6^) was performed using the FreeStyle MAX reagent. The transfection efficiency of these DNA constructs was determined by measuring the frequency of GFP^+^ cells using flow cytometry (Fig. EV5A). In certain assays, BMDCs (1 x 10^6^) were transfected (or not) with 20 μg of the neutralizing anti-IL-35 antibody (clone V1.4C4.22) using the Chariot protein delivery reagent (Active Motif). The efficiency of this antibody transfection was evaluated by subsequent (after 6 h) intracellular immunostaining of BMDCs with the secondary antibody Alexa Fluor 594-anti-mouse IgG and measuring the frequency of Alexa Fluor 594^+^ cells via flow cytometry (Fig. EV5B). To test whether the transfected anti-IL-35 antibody can bind intracellular IL-35, BMDCs were first transfected with the IL-35-expressing vector and then (after 24 h) with or without neutralizing anti-IL-35 antibody. Afterward, BMDCs were immunostained with the secondary antibody Alexa Fluor 594-anti-mouse IgG. The extent of anti-IL-35 antibody binding to overexpressed IL-35 was assessed by measuring GFP/Alexa Fluor 594 colocalization efficiency (yellow color after merging) using confocal microscopy (Fig. EV5C).

### ELISA

BMDCs (5 x 10^6^ cells/ml) were infected or not with LDPm for various times. In certain cases, BMDCs were washed and then treated with LPS (500 ng/ml) for 24 h. Culture supernatants were analyzed using ELISA kits specific for IL-12p70 (555256; BD Biosciences), TNFα (88-7324-88; Invitrogen) and IL-35 heterodimer (440507; BioLegend).

### Adoptive transfer of DCs

In this study, three adoptive transfer experiments were performed with BALB/c BMDCs to investigate whether different treatments or modifications of these BMDCs (see below) affect anti-leishmanial immune responses and parasite burden in LD-infected BALB/c mice after transfer. These modifications or treatments included silencing or inhibition of STAT3 with siRNA or WP1066 in BMDCs, respectively; overexpression of IL-35 in the STAT3-silenced or STAT3-inhibited BMDCs in some experimental sets; and transfection of BMDCs with a neutralizing anti-IL-35 or an isotype control antibody.

In the first adoptive transfer experiment, BALB/c BMDCs (1 x 10^6^) were transfected with control siRNA or STAT3-specific siRNAs. In some experimental groups, STAT3-silenced BMDCs were transfected with empty vector or IL-35-expressing vector. These BMDCs were subsequently administered intravenously to LD-infected BALB/c mice on days 15, 25, 35, 45 and 55 postinfection. Notably, repeated transfers of BMDCs were required to achieve a sustained and significant impact on anti-leishmanial immune responses, as our previous observations showed that adoptively transferred BMDCs persist in the spleens of recipient mice for only a short duration (up to 3 days) (Mishra *et al*., 2023). LD-infected mice that received or did not receive untreated DCs, as well as uninfected mice, were included as control groups for comparison.

In the second adoptive transfer experiment, BALB/c BMDCs (1 x 10^6^) were treated with WP1066 (5 μM) or DMSO (0.1%; control) for 1 h. As in the first experiment, some of the WP1066-treated BMDCs were transfected with IL-35-expressing plasmid or the corresponding empty plasmid. The BMDCs were then delivered intravenously to LD-infected BALB/c mice on the above-mentioned days postinfection. Again, the same sets of mice as in the first experiment were used as control groups.

In the third adoptive transfer experiment, BALB/c BMDCs (1 x 10^6^) were either left untransfected or transfected with an isotype control or a neutralizing anti-IL-35 antibody. BMDCs were then transferred to LD-infected BALB/c mice, as was done in the previous two adoptive transfer experiments. The control mice used here were the same as described above.

In all these experiments, the spleen and liver of all mice were isolated on day 60 after LD infection to measure the weight and parasite load of these organs. To determine the expression of IFNγ or IL-10 in CD4^+^ and CD8^+^ T cells, splenocytes from these mice were stained with PE/Cyanine7-anti-CD3 together with PerCP/Cyanine5.5-anti-CD4, or PE-anti-CD8 antibody. Subsequently, intracellular immunostaining was performed with FITC-anti-IFNγ or APC-anti-IL-10 antibodies using a Fixation/Permeabilization Buffer kit. The frequency of CD4^+^ and CD8^+^ T cells expressing IFNγ or IL-10 was then determined by flow cytometry. For adoptive transfer experiments involving anti-IL-35 antibody-transfected BMDCs, the frequency of the IL-35-expressing T cell population in the spleen of the above-mentioned mice was also assessed by flow cytometry, as described above.

### Treatment of LD-infected mice with neutralizing anti-IL-35 antibody

LD-infected BALB/c mice were injected intraperitoneally with a first dose (100 μg/mouse) of neutralizing anti-IL-35 antibody or isotype control antibody on day 15 post-LD infection. Subsequently, the anti-IL-35 antibody or the isotype control antibody was administered at a dose of 50 μg/mouse on days 25, 35, 45 and 55 post-LD infection. Age-matched uninfected mice and LD-infected mice that did not receive antibody treatment served as experimental controls. On day 60 postinfection, the weight and parasite load of the spleen and liver of these mice were determined. Additionally, the frequency of IFNγ- or IL-10-producing CD4^+^ and CD8^+^ T cells in the spleen was determined by flow cytometry.

### *In vivo* WP1066 treatment of LD-infected mice

Lyophilized WP1066 was reconstituted in a solvent mixture (vehicle) consisting of 80 parts polyethylene glycol (PEG) 300 and 20 parts DMSO (Sigma-Aldrich)(Kong *et al*, 2008) and administered to LD-infected mice via oral gavage at a dose of 20 mg/kg body weight, starting on day 15 postinfection. Subsequently, the same dose of WP1066 was administered orally three times a week (i.e. on alternate days) until day 54 postinfection. The control mice received either the vehicle (PEG 300/DMSO) or no oral gavage. After 60 days of infection, the weight and parasite load of the spleens and livers were measured.

### Assessment of *in vivo* parasite load

The livers and spleens of LD-infected mice were isolated to assess the *in vivo* parasite load. A portion (∼20 mg) of the spleen or liver was crushed to prepare a single cell suspension in RPMI-1640 complete medium, which was subsequently depleted of red blood cells. Genomic DNA was extracted from 2 x 10^6^ liver or spleen cells using the DNeasy Blood and Tissue Kit (Qiagen, Germany) and resuspended in 100 μl of nuclease-free water. DNA concentration was determined using a NanoDrop spectrophotometer. The qPCR reaction for each sample was carried out with 1 µl of DNA (nuclease-free water was used as a no-template control) and the primers specific for LD kinetoplast minicircle DNA [table EV1; (Rahim *et al*, 2022)] using the GoTaq qPCR kit (Promega). Parasite load/µl DNA was quantified by interpolation from a standard curve generated with a 10-fold serial dilution of the LDPm number from 10^5^ to 10^-1^, as described previously (Rahim *et al*., 2022). These data were used to measure the parasite load in 100 µl of DNA (obtained from 2 x 10^6^ spleen or liver cells). The parasite load in spleen and liver was finally expressed as Leishman-Donovan units (LDU), which was calculated as follows: Number of amastigotes per 1000 nucleated cells x liver or spleen weight (g) (Moreira *et al*, 2012).

### Flow cytometry

Flow cytometric analyses were performed using a FACSVerse and FACSAria flow cytometers (BD Biosciences). Data were analyzed using FlowJo software (Tree Star).

### Statistical analysis

One-way ANOVA (SigmaPlot 11.0 program) was used for all statistical analyses. A *P*-value <0.05 was considered statistically significant.

## Acknowledgments

The LD strain AG83 [MHOM/IN/83/AG83; American Type Culture Collection (ATCC PRA-413)] was obtained as a gift from Prof. Nahid Ali (CSIR-Indian Institute of Chemical Biology, India). IL-35 plasmid (pIGneo-mouse IL-35-GFP) and control plasmid (pIGneo-GFP) were generously given by Prof. Dario A. Vignali (University of Pittsburgh, USA). The pBabe vector containing mouse STAT3 and the empty control vector were kindly provided by Prof. Albert Baldwin (University of North Carolina at Chapel Hill, USA). HEK293T cells (CRL-3216; ATCC) were generously provided by Dr. Ashwani Kumar (CSIR-Institute of Microbial Technology, Chandigarh, India). We thank CSIR-IMTECH for providing research infrastructure, and CSIR-IMTECH animal house facility for providing mice required for experiments.

## Funding

This study was supported by grants from the Science and Engineering Research Board, Anusandhan National Research Foundation, Department of Science and Technology, India (CRG/2022/000292) and Council of Scientific and Industrial Research (OLP-0164 to P.S.). S.K. was supported by fellowships from University Grant Commission (UGC) and Science and Engineering Research Board, India. The funders had no role in study design, data collection and interpretation, or the decision to submit the work for publication.

## Contributions

S.K.: Conceptualization, methodology, investigation, validation, formal analysis and analyzed the data. Shubham: Investigation. M. M.: Investigation. R. K.: Investigation. P. S.: Conceptualization, methodology, validation, analysis, funding acquisition, resources, project administration, supervision, writing-original draft, writing-review and editing.

## Ethics declarations

The authors declare that they have no competing interests.

## Data availability

All relevant data are within the paper, Supplemental Material and Supporting Data files.

## Notes

### Competing Interest Statement

The authors have declared no competing interest.

